# Biodiversity and riparian forests are mutual biological drivers of ecosystem functions in temperate and tropical streams

**DOI:** 10.1101/2024.08.16.608233

**Authors:** Rebecca Oester, Paula M. de Omena, Larissa Corteletti da Costa, Marcelo S. Moretti, Florian Altermatt, Andreas Bruder

## Abstract

Fluxes of energy, matter, and organisms sustain linkages and functions within and between ecosystems. Yet, how biological drivers influence interactions and functions at the interface between aquatic and terrestrial environments (i.e., aquatic-terrestrial ecosystem functions) locally and across regions has received little attention. To test the relative importance of biological drivers on multiple aquatic-terrestrial ecosystem functions, we subsidised local terrestrial detritus in forested and non-forested stream sites in a temperate and tropical region. We also manipulated leaf litter diversity (horizontal biodiversity of resources) and macroinvertebrate access (vertical biodiversity of consumers). We measured secondary production of aquatic fungi, in-stream leaf litter nitrogen loss, and decomposition rates. The simultaneous provision of all three ecosystem functions (i.e., multifunctionality) was positively driven by vertical biodiversity and riparian forests in both regions. In both tropical and temperate streams, nitrogen loss was associated with vertical biodiversity. Decomposition rates were also enhanced by vertical biodiversity and linked to other ecosystem functions. These results reveal strong and consistent effects of biodiversity and riparian forests on aquatic-terrestrial ecosystem functions in freshwater detrital food webs in both temperate and tropical headwater streams. Thus, disentangling the drivers of ecosystem functions in these systems requires an understanding of underlying mechanisms beyond ecosystem borders.

## Introduction

Between adjacent ecosystems, fluxes of energy, matter and organisms create tight linkages (Barnes *et al*. 2018), often promoting biodiversity (Kark 2013) and enhancing ecosystem processes (Naiman *et al*. 1993; Tolkkinen *et al*. 2020). For example, subsidy flows across ecosystem boundaries, not only foster complex trophic interactions reliant on both ecosystems but also support biodiversity at their interfaces (Baxter *et al*. 2005). Many trophic interactions between resources and consumers from riparian and freshwater ecosystems are sustained by subsidies from the riparian vegetation into aquatic food webs (Ferreira *et al*. 2023) in line with the river-continuum concept (Vannote *et al*. 1980), meta-ecosystem theory (Gounand *et al*. 2018; Harvey *et al*. 2023), and empirical studies (e.g., Rezende *et al*. 2019; Webster & Meyer 1997).

Subsidies in the form of allochthonous leaf litter provide the main resource for various decomposers and detritivores in forested streams (Cummins *et al*. 1989; Wantzen & Wagner 2006). As leaf litter enters the streams, microbes, such as aquatic fungi, are the first to colonise it (Gessner & Chauvet 1997). Their activity and growth can change leaf litter stoichiometry through nutrient immobilization and mineralization (García-Palacios *et al*. 2017). Detritivores, together with microbial decomposers, control decomposition rates and nutrient release of leaf litter (Gessner *et al*. 2010; Handa *et al*. 2014). The diversity in leaf litter species (horizontal biodiversity of resources) can enhance niche complementarity in detrital food webs (Loreau & Hector 2001), and increase decomposer and detritivore diversity (vertical biodiversity of consumers; Gessner *et al*. 2010; Scherer-Lorenzen *et al*. 2022). As a result, higher resource and consumer diversity together can enhance ecosystem functioning (Figure 1A; Duarte *et al*. 2006; Jabiol *et al*. 2013; Kominoski *et al*. 2010; Liu *et al*. 2020). Key ecosystem functions in freshwater detrital food webs are the processing, nutrient dynamics, and subsequent transformation of terrestrial detritus into secondary production (Dang *et al*. 2005; Dodds *et al*. 2004; Duarte *et al*. 2006; Gessner & Chauvet 1997). Thus, cross-ecosystem linkages and diversity of resources and consumers can benefit ecosystem functions across ecosystem boundaries (Scherer-Lorenzen *et al*. 2022).

**Figure 1:**
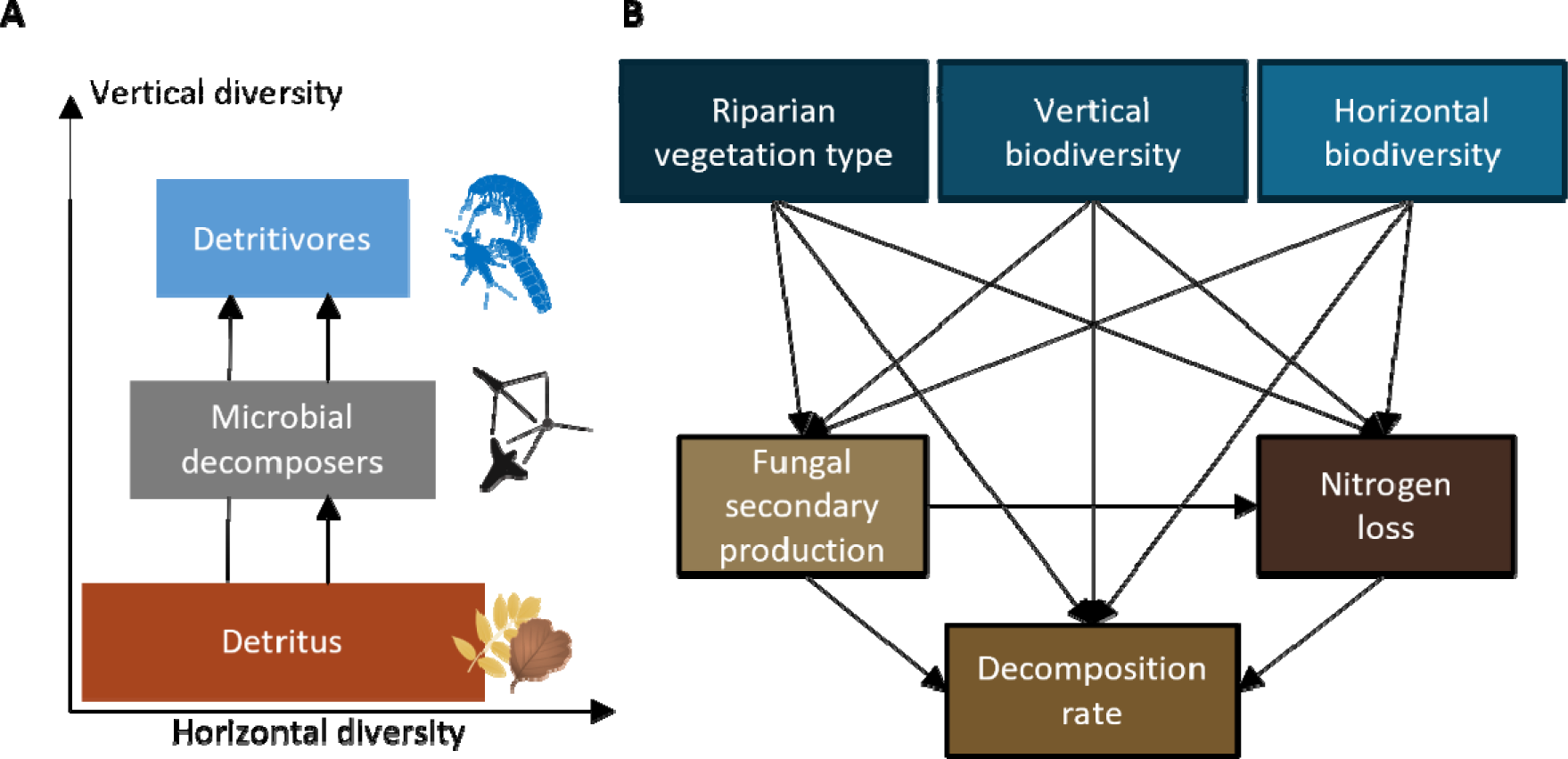
Conceptual figure of detrital food webs in headwater streams and links influencing ecosystem functions. A: Detrital food web components represented in horizontal diversity (biodiversity within trophic guild) and vertical diversity (biodiversity between trophic guilds; modified from Gessner et al. 2010). B: Meta-model on the direct and indirect effects on ecosystem functions represented as path diagram. Detailed hypotheses and literature references for each link can be found in Table S1.

However, land-cover alterations (Little & Altermatt 2018) and biodiversity decline (López-Rojo *et al*. 2019) affect terrestrial subsidies in freshwaters. For instance, land cover changes affecting the riparian forest also influence adjacent aquatic ecosystems (Tolkkinen *et al*. 2020) not only abiotically (Ferreira *et al*. 2023; Naiman *et al*. 1993) but also through changes in quantity, diversity, and composition of leaf litter input (Ohler *et al*. 2023). These examples illustrate how human impacts can cascade across ecosystems and affect biodiversity, cross-ecosystem linkages and ecosystem functions (Handa *et al*. 2014). Understanding, conserving and maintaining these subsidy flows are not only essential for biodiversity (Naiman *et al*. 1993; Scherer-Lorenzen *et al*. 2022), and food-web dynamics (Barnes *et al*. 2018; Beiser *et al*. 1991; Jabiol *et al*. 2013), but also the capability of streams to simultaneously perform multiple ecosystem functions (i.e., multifunctionality; Ferreira *et al*. 2023; Lefcheck *et al*. 2015; Soliveres *et al*. 2016).

Based on the growing literature on the relationship between biodiversity and ecosystem functions (i.e., BEF framework; Lefcheck *et al*. 2015; Soliveres *et al*. 2016), the degree of multifunctionality depends on the dominant mechanisms and relationships between each ecosystem function considered (Turnbull *et al*. 2016). In ecosystems with high biodiversity and ecological complexity, such as forested headwater streams, we might expect complementarity effects to outweigh other mechanisms (e.g., interference competition or resource elimination). Complementarity effects would result in overall positive biodiversity effects on ecosystem functions (Kominoski *et al*. 2010; Loreau & Hector 2001). However, an ecosystem function is often controlled by multiple environmental conditions simultaneously and by feedbacks from other ecosystem functions (Eisenhauer *et al*. 2019; Soliveres *et al*. 2016), contributing to the complexity of BEF-relationships. Thus, ecosystem multifunctionality might ultimately seem to be context-dependent (Catford *et al*. 2022). For example, the importance of consumer diversity on decomposition rates might be higher in the tropics compared to temperate regions (Boyero *et al*. 2021). In contrast, resource characteristics might be the dominant drivers in temperate regions (López-Rojo *et al*. 2019). Uncertainty about the generality of these drivers of ecosystem functions across multiple biomes still challenges predictions of multifunctionality in natural settings (van der Plas 2019; Wantzen & Wagner 2006).

To investigate the relative importance of biological drivers of aquatic-terrestrial ecosystem functions (i.e., functions that occur at the interface and combine both components of aquatic and terrestrial environments), we tested the effects of resource and consumer biodiversity, and riparian forest presence on multiple ecosystem functions in detritus-based food webs in temperate and tropical streams (Figure 1B). We performed replicate field experiments in headwater streams in Switzerland and Brazil (Figure 2) as two regions with distinct climates (Shah *et al*. 2017), detritus quality (Boyero *et al*. 2017), and aquatic biodiversity (Boyero *et al*. 2021; Wantzen & Wagner 2006). We used standardised leaf litter decomposition assays to test how riparian forest (non-forested vs. forested), horizontal biodiversity of resources (single leaf species vs. two leaf species mixed), and vertical biodiversity of decomposer community (microbial only vs. microbial + macroinvertebrate) influence the individual and combined (multifunctionality) aquatic-terrestrial ecosystem functions: secondary production of aquatic fungi growing on leaf litter, nutrient dynamics measured as in-stream nitrogen loss from leaf litter, and aquatic leaf litter decomposition rates (Dang *et al*. 2005; Duarte *et al*. 2006). We expected large environmental differences to influence the relative importance of these different drivers of ecosystem functions (Boyero *et al*. 2021; Handa *et al*. 2014; Soliveres *et al*. 2016). Yet, we predicted overall positive effects of riparian forests, vertical, and horizontal biodiversity on ecosystem functions in both regions due to beneficial effects of completeness of decomposer community (complementarity effects) but with litter species-specific mixing effects (Frainer *et al*. 2015; Handa *et al*. 2014; Schindler & Gessner 2009).

**Figure 2:**
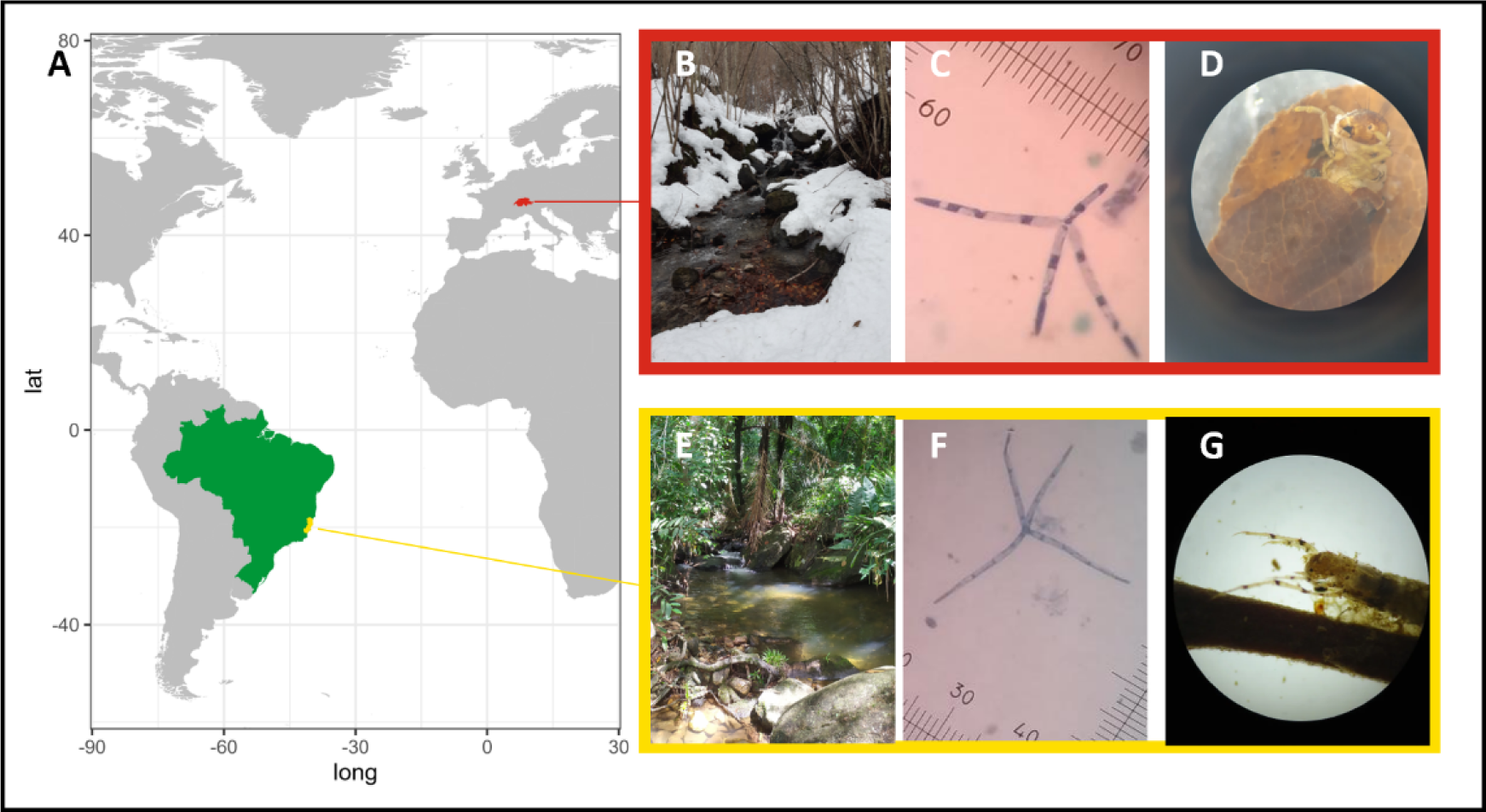
Map and pictures of study sites and local detritus-associated biodiversity. A: Temperate and tropical study regions highlighted in red (Switzerland) and green (Brazil) with the state of Espirito Santo highlighted in yellow. B and E: Study sites in each region during the field experiment. C and F: Stained conidia of aquatic fungi *Articulospora tetracladia* (photo: J. Jabiol), and *Lemmoniera aquatica* (photo: J. Jabiol), respectively. D and G: Caddisflies (Trichoptera) from Switzerland and Brazil with cases made of detritus: Limnephilidae and Leptoceridae detritivores, respectively.

## Material and Methods

### Experimental design

To test the local effects of riparian forests on ecosystem functions in temperate and tropical regions, we ran standardised leaf litter decomposition experiments in headwater streams in Switzerland and Brazil (Atlantic Forest biome in the state of Espirito Santo, SE), always contrasting forested and non-forested stream sections (Figure 2). We selected eight streams in each country, each having a distinct section with the riparian vegetation consisting of densely standing trees (forested) and another section surrounded by locally typical grassland or extensively used pastures with no or only isolated trees (non-forested). This contrast of forested vs. non-forested stream sections is a commonly found landscape pattern of the studied stream systems (Casotti *et al*. 2015; Cereghetti & Altermatt 2023; Little & Altermatt 2018). The forested site was upstream in half of the streams and downstream in the other half, which resulted in a balanced and paired comparison between forested and non-forested sites within each stream. The sites were located at elevation levels where the natural riparian vegetation would consist of deciduous forests. Other than the forest-non-forest transition, there were only minimal anthropogenic disturbances or modifications, thus similar nutrient and temperature levels within the streams (for details on stream and riparian zone characteristics see Table S2).

In each site, we tested the effects of horizontal biodiversity of resources (i.e., leaf litter diversity) by mixing leaf litter from a local nitrogen-fixing tree species and a species rich in labile carbon in mesh bags. We used 5 g (SD = 0.03 g) of dried leaf litter for each leaf litter bag. Thus, the leaf litter consisted of either a single species (mono) or two species (mixed in equal mass) of the local species pool. In Switzerland, these were *Alnus glutinosa* (L.) Gaertn. (Betulaceae) and *Fraxinus excelsior* (L.) (Oleaceae) and in Brazil *Inga laurina* (Sw.) Willd. (Fabaceae) and *Miconia chartacea* Triana (Melastomataceae), respectively. These four tree species are locally common riparian species, with relatively high to moderate nutritional quality and palatability for detritus consumers (Bruder *et al*. 2014; Kiffer *et al*. 2018; Schindler & Gessner 2009). Details on the initial leaf litter characteristics can be found in Table S3.

To test the effects of vertical biodiversity of consumers (i.e., trophic complexity), we used leaf litter bags with different mesh sizes in each site to be naturally colonised by local decomposers and detritivores. We used fine and coarse mesh bags (0.25 mm and 10 mm opening, respectively) to exclude or allow the access of macroinvertebrate detritivores. Both types of bags allowed for microbial colonization of the leaf litter. Each treatment combination (horizontal and vertical) was replicated four times, resulting in a total of 24 leaf litter bags per site (768 bags in total). This fully crossed, experimental design allowed us to assess the individual effects of riparian forests and horizontal and vertical biodiversity on multiple ecosystem functions associated with detrital food webs in streams.

We started the field experiments during the local leaf fall season and terminated them when approximately 50 % of the initial leaf litter mass was lost. Consequently, the duration and timing of the field experiment varied between regions, resulting in three to five weeks from December 2020 to January 2021 in Switzerland and six to eight weeks from May to July 2021 in Brazil. We excluded data from leaf litter bags with < 25 % leaf litter mass remaining (n = 5, i.e., 1.3 %), as consumer colonization and activity might be reduced (Beiser *et al*. 1991).

### Ecosystem functions

To assess leaf litter decomposition, we quantified the decomposition rate for each leaf litter type in each litter bag following the exponential decay model: m_t_ = m e ^-kt^, where m_t_ is the leaf litter dry weight after t degree days, m_0_ is the initial dry weight, and k is the decomposition rate (Webster & Benfield 1986). We calculated degree-days based on hourly measurements of stream temperature (HOBO Temperature DataLogger; UA-002-64; Onset Computer Corporation) and standardised process rates to degree-days to account for temperature-dependent consumer activity as similarly done in cross-biome studies (Bruder *et al*. 2014; Ferreira *et al*. 2019; Wantzen & Wagner 2006).

To determine fungal secondary production, we quantified fungal biomass in the leaf litter at the end of the experiment. We quantified ergosterol, a fungal cell membrane compound, from ten leaf disks cut from different leaves per litter bag and species. After subsampling, we stored the leaf disks at -20°C, then freeze-dried them and extracted and purified ergosterol with solid-phase extraction (Sep-Pak® Vac RC tC18 500 mg sorbent; Waters, Milford, USA; Gessner 2020). We then measured the ergosterol concentration using ultra-high-performance liquid chromatography (UHPLC; 1250 Infinity Series, Agilent Technologies, Santa Clara, USA) at a wavelength of 282 nm and estimated fungal biomass with a conversion factor of 5.5 mg ergosterol per g fungal biomass (Gessner 2020).

To quantify the detrital nutrient dynamics, we measured the N content of the leaves from dried (60 °C for 48 h), homogenised, and encapsulated (9 × 5 mm tin capsules, Säntis Analytical) leaf litter material (1–1.5 mg) on an Elemental Analyzer (Elementar, Vario Pyro). We calculated the nitrogen (N) loss (%) according to Handa *et al*. (2014): 100L×L[(M_i_L×LN_i_)L−L(M_f_L×LN_f_)]L/L(M_i_L×LN_i_), where M_i_ and M_f_ are the initial and final leaf litter dry mass, respectively, and N_i_ and N_f_ are the initial and final N concentration (% of leaf litter dry mass), respectively. Initial N_i_ was quantified from a representative subset of leaves from the original batch.

### Statistical analysis

We performed all calculations and statistical analyses in R (version 4.1.2; R Development Core Team, 2020). To assess multifunctionality, we log-transformed when not normally distributed, scaled (z-scored), and averaged data of the three ecosystem functions studied, i.e., fungal secondary production (biomass of aquatic fungi at the end of the experiment), N loss (% N lost from the leaves at the end of the experiment), and leaf decomposition rate (k) for each leaf litter species using the multifunc package (Byrnes 2022). The highest multifunctionality scores are achieved if leaves show high fungal secondary production (Gessner & Chauvet 1997), high N loss (García-Palacios *et al*. 2017; Handa *et al*. 2014), and high decomposition rates (Dang *et al*. 2005). As the averaging of functions inherently leads to a simplification, has some statistical limitations, and can suffer from biases in defining the process maxima and minima (Meyer *et al*. 2018), we also reported results for each function individually in the supplementary material (Figure S2, S3, Table S4, S5). We used the mean multifunctionality score as a response variable and tested the effects of riparian forest, horizontal and vertical diversity separately for the different leaf species in a Bayesian generalised non-linear multivariate multilevel models (brms; Bürkner *et al*. 2021), which use the probabilistic programming language for statistical interference “Stan” (Stan Development Team 2023). As the multifunctionality score lied between 0 and 1, we chose weakly informative priors of normally distributed intercepts and slopes (0.5, 1). To account for the study design and environmental differences among streams, we incorporated sampling location as random effects. For all models, we generated 4,000 (four chains run for 2,000 iterations discarding the first 1,000 as burn-in) Markov chain Monte Carlo (MCMC) samples from the posterior distribution where draws were sampled using NUTS (No-U-Turn Sampler).

To assess the detailed direct and indirect effects of riparian forest, horizontal and vertical biodiversity, over fungal secondary production, on N loss and leaf litter decomposition rates, we constructed path diagrams (i.e., structural equation models; SEMs) again using Bayesian generalised non-linear multivariate multilevel models (brms; Bückner 2019). Based on our hypotheses (Table S1) and the metamodel (Figure 1B), we used the same model structure but modelled each leaf species separately to assess the species-specific effects on ecosystem functions. We z-transformed all numeric values and included weakly informative priors of normally distributed intercepts (0, 1) and slopes (0, 0.5). To account for the study design and environmental differences among streams, we again incorporated sampling location as random effects. For all models, we generated 20,000 (four chains run for 10,000 iterations, discarding the first 5,000 as burn-in) MCMC samples from the posterior distribution where draws were sampled using NUTS. For all models, MCMC chains indicated convergence by falling within the threshold specified by Gelman & Rubin (1992) and showing high effective sample size measures. We visually checked the fit of the posterior distribution with the data (bayesplot, Gabry *et al*. 2021).

## Results

We found strong evidence for increased multifunctionality in sites with riparian forests and vertical biodiversity in both Swiss and Brazilian streams (Figure 3). In both temperate and tropical leaf species, scaled mean multifunctionality was statistically significantly higher in forested compared to non-forested sites (*Alnus*: estimate = 0.03, CIs = [0.01, 0.04]; *Fraxinus*: 0.03[0.01, 0.05]; *Inga*: 0.02[0.01, 0.03]; *Miconia*: 0.05[0.02, 0.07]). Moreover, for *Alnus*, *Fraxinus,* and *Miconia*, we also found higher multifunctionality scores with the inclusion of higher trophic level consumers (*Alnus*: 0.10[0.08, 0.12]; *Fraxinus*: 0.12[0.10, 0.14]; *Miconia*: 0.12[0.10, 0.14) but not for *Inga* (-0.00[-0.02, 0.01]). Across all leaf species, multifunctionality scores consistently showed no influence of horizontal diversity of leaf litter (*Alnus*: -0.02[-0.04, 0.00]; *Fraxinus*: -0.00[-0.02, 0.02]; *Inga*: 0.01[-0.01, 0.02]; *Miconia*: 0.02[-0.00,0.04]).

**Figure 3:**
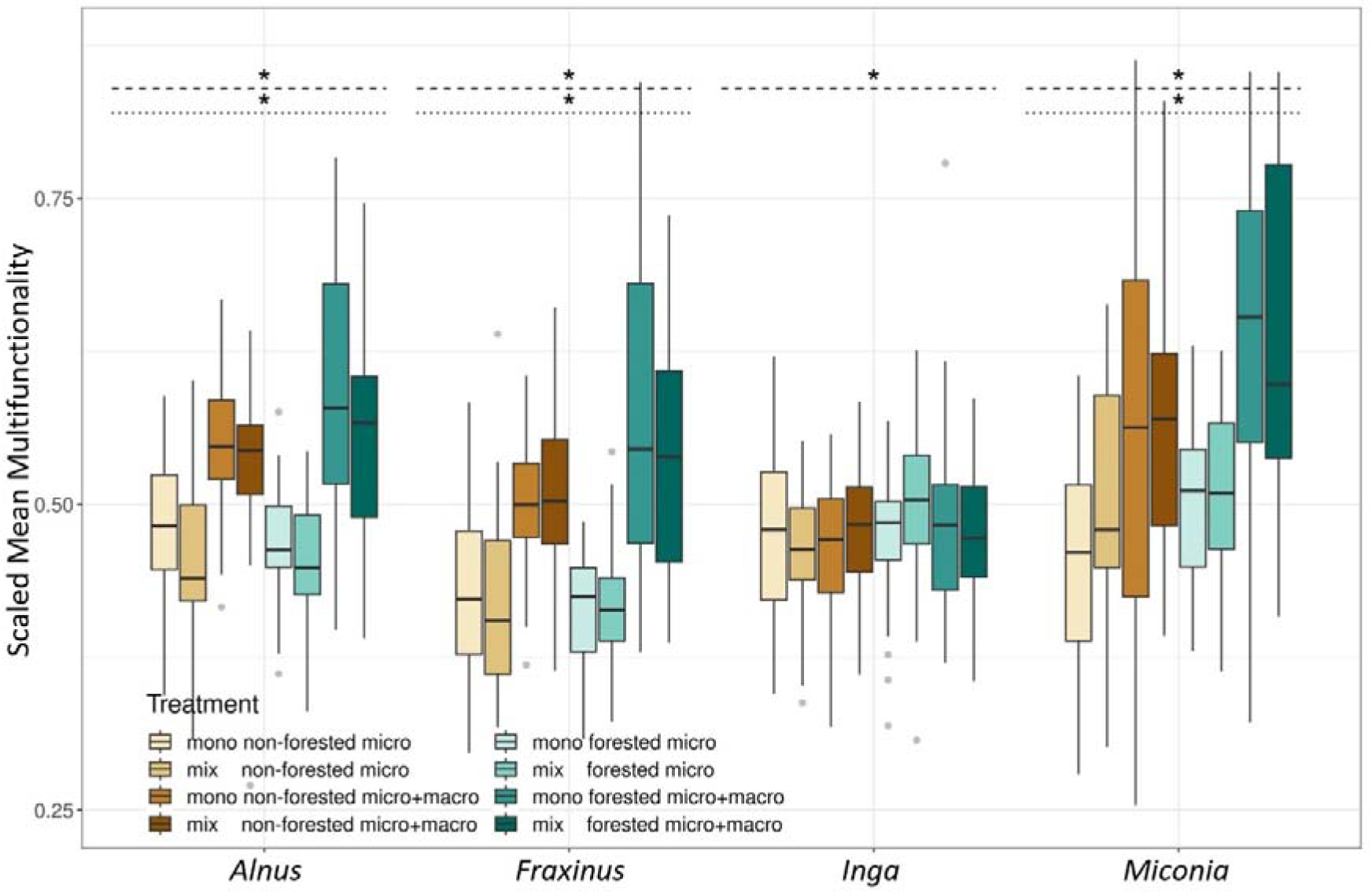
Boxplots of scaled mean multifunctionality depending on leaf species and experimental treatment. Colours differentiate the treatment combinations of riparian forest and horizontal and vertical biodiversity, with mixed (mix) or single (mono) species leaf litter bags, placement in a forested or non-forested stream section and inclusion (micro+macro) or exclusion (micro) of macroinvertebrate consumers. Dashed lines indicate the effects of riparian vegetation type (forested vs. non-forested) and dotted lines show the effects of vertical biodiversity (micro vs. micro+macro) separately assessed for each leaf species.

When assessing the individual effects of biodiversity and riparian forests on stream ecosystem functions related to detritus, our results showed that almost all effects of riparian forests and biodiversity are positive (shown as blue arrows in Figure 4) and consistent between most leaf litter species (shown as medium and thick arrows in Figure 4). Leaf litter in forested compared to non-forested sites showed consistently higher nitrogen (N) loss for all four leaf species (c in Figure 4). We found on average 12, 18, 22, and 273 % increase in N loss comparing forested and non-forested sites for *Inga*, *Ash*, *Alder,* and *Miconia*, respectively (Table S7). For *Alnus*, *Fraxinus,* and *Inga,* we additionally found positive, yet not statistically significant effects of riparian forests on decomposition rates (b in Figure 4), and for *Miconia* we also found positive effects of riparian forests on fungal secondary production (a in Figure 4).

**Figure 4:**
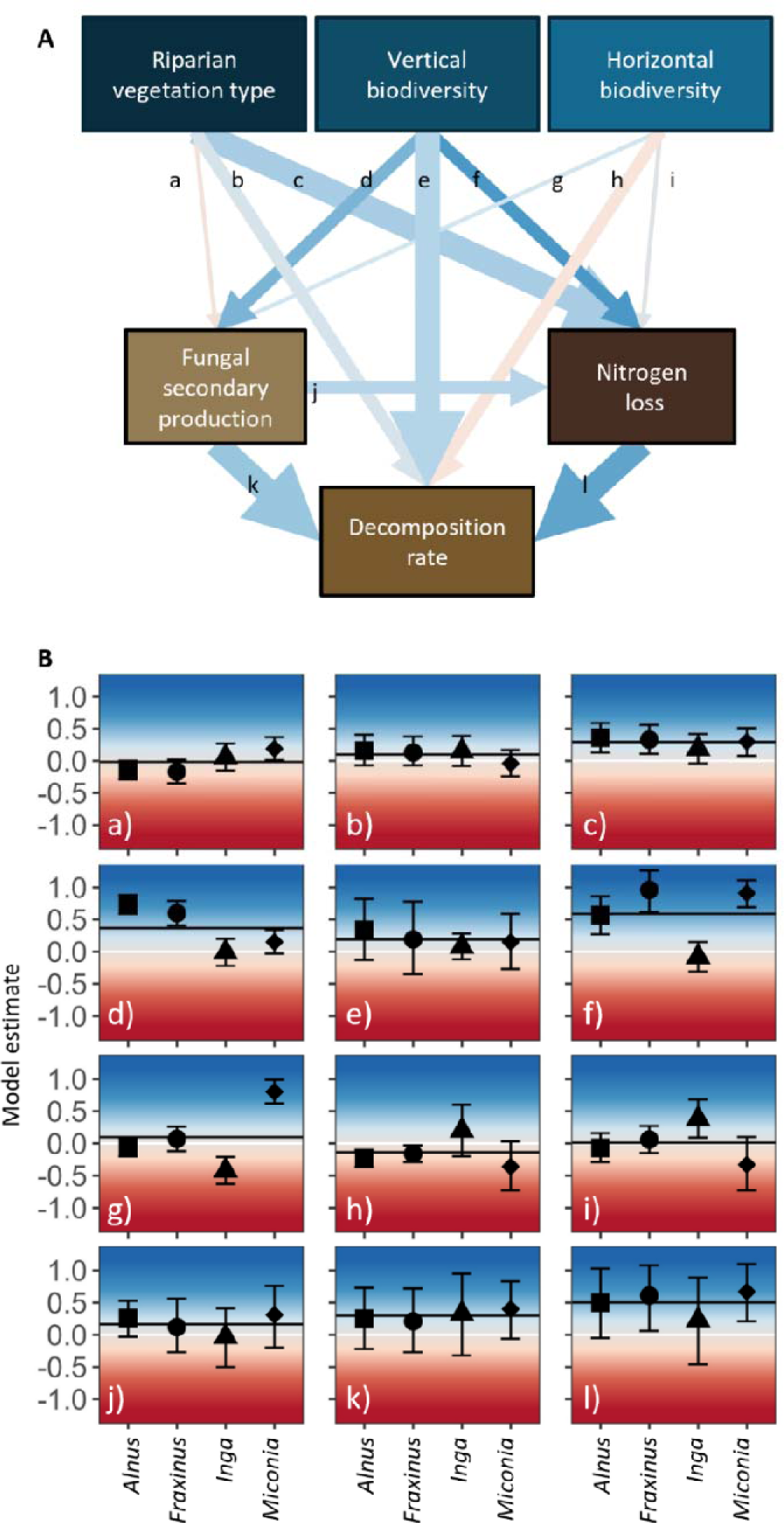
Path model (SEM) and its effect sizes linking biological drivers (blue boxes) to individual ecosystem functions (brown boxes). A: Effects of biological drivers are always contrasting non-forested vs. forested vegetation; micro vs. micro+macro community; and mono vs. mixed leaf species. The colour of the arrows displays the average effect size ranging from blue (highly positive) to red (highly negative) consistent with the colour palette in B. Thick arrows show consistency in effect direction for all four leaf species, and normal arrows show that three of the four leaf species show consistent directionality, while fine arrows show inconsistent directionality. B: Model estimates of the SEMs for each leaf species and separated by each path with the letters separating each panel representing the paths labelled in panel A. The error bars represent CIs. The horizontal white line indicates 0 and the horizontal black line indicates the overall mean effect size across all four leaf species.

Vertical biodiversity affected all three ecosystem functions positively yet depending on the leaf species to different extents. The macroinvertebrate access (micro + macro) compared to their exclusion (micro) to the leaf litter bags increased fungal secondary production on *Alnus* and *Fraxinus* significantly but had little effects on *Inga* and *Miconia* (d in Figure 4). Our data also suggested consistent but statistically insignificant positive effects of vertical biodiversity on decomposition rates of all four leaf species (e in Figure 4). We also found significantly positive effects of macroinvertebrates on N loss for *Alnus*, *Fraxinus* and *Miconia*, but not *Inga* (f in Figure 4).

Horizontal biodiversity had mostly species-specific effects as mixing different leaf species resulted in both positive and negative effects for all three ecosystem functions (g, h, i in Figure 4). In particular, *Miconia* showed higher fungal secondary production when together with *Inga*, while the opposite pattern occurred for *Inga* when together with *Miconia* (g in Figure 4). For *Alnus* and *Fraxinus*, no such mixing effect occurred for fungal secondary production. Still, both leaf species decomposed slower in mixed leaf litter bags (h in Figure 4).

We also found consistently positive effects among ecosystem functions. Fungal secondary production, N loss, and decomposition rates in all leaf litter species were positively correlated. However, these relationships compared to treatment effects, showed large CIs and were not always statistically significant (j, k, l in Figure 4, Figure S3).

## Discussion

Ecosystem functions in detrital stream food webs were overall positively driven by vertical biodiversity (trophic diversity) and the presence of riparian forests around both temperate and tropical headwater streams. Predominantly, we observed high multifunctionality scores in the most complex food web configurations and forested stream sections. Specifically, nitrogen (N) loss and decomposition rates were positively influenced by the presence of riparian forests and by vertical biodiversity. However, horizontal biodiversity (resource diversity) had species-specific impacts on different ecosystem functions. In both temperate and tropical streams, ecosystem functions were similarly associated with each other. These overall positive responses likely reflect complementarity effects outweighing other BEF-mechanisms locally, as well as underlying similarities in how these temperate and tropical stream food webs function. Our findings thus identify and highlight biodiversity and riparian forests as important biological drivers of aquatic-terrestrial ecosystem functions in freshwater detrital food webs. These insights also emphasise the similarities in cross-ecosystem linkages in environmentally distinct biomes. Given the scale and speed of current land-cover changes, further understanding, conserving, and maintaining these cross-ecosystem linkages and drivers are essential not only for aquatic communities and their food web dynamics, but also the many ecosystem functions they provide.

Independent of biogeographic region, multifunctionality in detrital stream food webs was positively influenced by riparian forests and vertical biodiversity (Figure 3). In other words, the capacity of the studied headwater streams to perform multiple ecosystem functions depended on the presence of riparian forests and the trophic completeness of the aquatic detrital food web. These results highlight that biodiversity at multiple trophic levels benefits ecosystem multifunctionality (Soliveres *et al*. 2016) also in headwater streams. The overall positive effects of riparian forests and biodiversity on aquatic-terrestrial ecosystem functions in the temperate and tropical freshwater detrital food web may point to fundamental underlying mechanisms determining how these ecosystems function. First, complementarity effects (caused by facilitation and resource partitioning) resulting in increased total resource use can enhance ecosystem functioning in diverse communities through positive interactions among species (Loreau & Hector 2001). This BEF-effect has been demonstrated numerous times in terrestrial ecosystems (e.g., Loreau & Hector 2001), but also in detritus-based streams (Handa *et al*. 2014; Jabiol *et al*. 2013; Kominoski *et al*. 2010). Second, while temperate and tropical streams differ in many aspects (Boyero *et al*. 2017, 2021; Shah *et al*. 2017), microbial colonization and growth on riparian leaf litter, detritivore feeding, and subsequent nutrient and decomposition dynamics follow the same principles and can be subject to complementary effects (Bruder *et al*. 2014; Ferreira *et al*. 2019; Wantzen & Wagner 2006). While we showed these food-web responses based on a fully factorial two-level design of three relevant biological drivers on three major ecosystem functions in headwater streams (Dang *et al*. 2005; Duarte *et al*. 2006), the inclusion of continuous multitrophic biodiversity (Barnes *et al*. 2018; Eisenhauer *et al*. 2019; Soliveres *et al*. 2016) in addition to environmental gradients might result in even finer-scaled differentiation and “biodiversity fingerprint” in future studies (Lefcheck *et al*. 2015). As replicated field experiments across biomes are still rare (but see e.g., Boyero *et al*. 2021; Bruder *et al*. 2014; Ferreira *et al*. 2019; Handa *et al*. 2014; Wantzen & Wagner 2006), our results provide a first cross-biome comparison of drivers of ecosystem multifunctionality in freshwater detrital food webs.

Our results suggest many similarities in how riparian forests and biodiversity drive different aquatic-terrestrial ecosystem functions (Figure 4). Besides the important effects of riparian forests on abiotic conditions of stream ecosystems (e.g., regulation of microclimate, erosion, and hydrology; Ferreira *et al*. 2023; Tolkkinen *et al*. 2020), our findings highlight and quantify the essential role of riparian forests for the composition and quantity of subsidies in the form of leaf litter for freshwater detrital food webs and their functions (Gounand *et al*. 2018; Vannote *et al*. 1980). Notably, riparian forests significantly benefitted detritus-based nutrient cycling through increased N turnover. Possible explanations could lie in the more suitable environmental and resource conditions for consumers (Tolkkinen *et al*. 2020), increasing their diversity, abundance, biomass (Oester *et al*. 2023), and overall leaf litter consumption rate (Oester *et al*. 2024 *in press*). Since N is only net released (mineralization > immobilization) in the later stages of leaf decay (García-Palacios *et al*. 2017), a faster turnover indicates a higher nutrient release from the resources to the environment. This released N can be used by consumers or get reabsorbed by the riparian vegetation.

Another similarity was reflected in the predominantly positive effects of vertical biodiversity on aquatic-terrestrial ecosystem functions of both temperate and tropical freshwater detrital food webs. While these results are not surprising, as aquatic detritivores rely mostly on terrestrial resources (Danger *et al*. 2012), these insights emphasise the strong influence of macroinvertebrate detritivores on multiple ecosystem functions in both regions. While detritivores fragment and feed on leaf litter often already colonised by microbes (Danger *et al*. 2012; Duarte *et al*. 2006), they can also affect local nutrient levels through bioturbation and excretion (Chakraborty *et al*. 2022; Wallace & Webster 1996). However, their importance for these ecosystem functions might depend on the local biodiversity and abundances, especially in less species-rich biomes (Boyero *et al*. 2021; Bruder *et al*. 2014; Handa *et al*. 2014; Wantzen & Wagner 2006).

Generally, a more complex trophic network with more trophic levels not only increases the potential for multiple ecosystem functions but also buffers them against disturbances (Jabiol *et al*. 2013; Soliveres *et al*. 2016). However, only the increased vertical but not horizontal food web complexity led to consistent increases in ecosystem functions. As expected, the consequences of leaf litter mixing depended on the functional composition of the leaf litter, likely influenced by differing stoichiometry and other chemical compounds (Schindler & Gessner 2009; but see Frainer *et al*. 2015). Although all four leaf species used have relatively labile leaf litter (Bruder et al. 2014; Kiffer et al. 2018; Schindler & Gessner 2009), they vary in chemical (Table S3) and physical characteristics, such as toughness (Bruder *et al*. 2014; Moretti *et al*. 2007; Rezende *et al*. 2019). For example, *Inga* showed the lowest values in several ecosystem functions (Table S7), likely due to the less advanced processing stage, as *Inga* leaf litter was the least nutritious (lowest N content) and hardest to decompose (highest amount of lignin; Table S3). Hence, mixing *Inga* with *Miconia* and *Alnus* with *Fraxinus* lead to varying results for all three ecosystem functions. This leaf litter mixing effect might have also been mediated by resident macroinvertebrates as shown for temperate (Santonja *et al*. 2020) and tropical ecosystems (Rabelo *et al*. 2024), as well as in global assessments (Handa *et al*. 2014; Liu *et al*. 2020). As riparian forests provide a variety of leaf litter (Little & Altermatt 2018; Wantzen & Wagner 2006), especially in the tropics (Boyero *et al*. 2017; Rabelo *et al*. 2024), it is important to assess the consequences of leaf litter mixing on multiple ecosystem processes to better understand the complexity of in-stream conditions and functions.

Despite major environmental differences between temperate and tropical stream ecosystems, we found several mutual biological drivers of multiple ecosystem functions in their detrital food webs. Our study thus corroborates earlier ones in that biodiversity at multiple trophic levels as well as riparian forests are essential for stream ecosystems to fulfil their various functions. Thus, to understand the factors influencing ecosystem functions, it is essential to comprehend the underlying processes and mechanisms that extend beyond ecosystem boundaries. Ultimately, a better understanding of the ecological potential of small streams and their riparian vegetation as hotspots for biodiversity and ecosystem functions is needed to respond to the challenges posed by global change and biodiversity decline.

### Use of Artificial Intelligence (AI) and AI-assisted technologies

We have not used AI-assisted technologies in creating this article.

## Author contribution

Conceptualization: RO, PO, LC, MM, FA, AB; Methodology: RO, PO, LC, AB; Writing: RO, PO, LC, MM, AB, FA; Funding acquisition: AB, MM, FA; Supervision: AB, MM, FA

## Data availability

The data and code that support the findings of this study are openly available on Dryad, Zenodo, and GitHub: https://doi.org/10.5061/dryad.r7sqv9smr; https://github.com/RebeccaOester/Biodiversity-and-riparian-forests-are-mutual-biological-drivers-of-ecosystem-functions-in-streams

## Acknowledgment

We thank Eva Cereghetti, Francesca Cerroti, Lars Sturm, Ali Reza Esmaeili, Lyandra Oliveira, Marcos Ferraz and Karoline Serpa for their help with fieldwork, and all landowners whose property we crossed to access our sampling sites. We are also grateful for N. Dubois and I. Brunner for supporting the stoichiometric analyses. Lastly, we thank anonymous reviewers for comments on a previous version of the manuscript and the funding agencies SNF, URPP, FAPES and CNPq.

