## Supplementary Material for "Biodiversity and riparian forests are mutual biological drivers of ecosystem functions in temperate and tropical streams"

**Affiliations:**
1 Institute of Microbiology, University of Applied Sciences and Arts of Southern Switzerland, via Flora Ruchat Roncati 15, CH-6850 Mendrisio, Switzerland

2 Department of Evolutionary Biology and Environmental Studies, University of Zurich, Winterthurerstr. 190, CH-8057 Zurich, Switzerland

3 Eawag: Swiss Federal Institute of Aquatic Science and Technology, Department of Aquatic Ecology, Überlandstrasse 133, CH-8600 Dübendorf, Switzerland

4 Laboratory of Aquatic Insect Ecology, University of Vila Velha, Av. Comissário José Dantas de Melo 21, 29102-920 Vila Velha, ES, Brazil

**Table S1**: Hypotheses for the construction of the SEM meta-model. Each link between predictor and response variable is based on hypotheses derived from existing literature on temperate and/or tropical streams. The hypotheses represent either a positive (+), negative (-) or neutral (0) effect. Hypotheses for the drivers: riparian vegetation type (forested vs. non-forested), vertical biodiversity (micro + macro vs. micro only), and horizontal biodiversity (two leaf litter species in a mixture vs. single leaf litter species) are based on the difference between the reference state (non-forested, micro, single leaf species) and the more ecologically complex state (forested, micro+macro, mixed leaf species). For example, we expect higher fungal secondary production in a forested site compared to a non-forested site, represented as a (+). Fungal secondary production is measured as fungal biomass in the remaining leaf litter, N loss is measured as the % N released from the leaf litter, and decomposition rate is measured as k based on an exponential decay function described in the main text. The letters A-L correspond to Figure 4 in the main text.

| Variables | Link from | Ecosystem functions | Letter | H_1_ CH | H_1_  BR | Sources |
| --- | --- | --- | --- | --- | --- | --- |
| exogeneous | Riparian  Vegetation type | Fungal secondary production | A | 0 | 0/+ | (1, 2) |
|  | Riparian  Vegetation type | Decomposition rate | B | + | + | (3–5) |
|  | Riparian  Vegetation type | N loss | C | + | + | (6–9) |
|  | Vertical  Biodiversity | Fungal secondary production | D | 0/- | 0 | (10–12) |
|  | Vertical  Biodiversity | Decomposition rate | E | + | 0/+ | (13, 14) |
|  | Vertical  Biodiversity | N loss | F | + | 0 | (12, 15–18) |
|  | Horizontal  Biodiversity | Fungal secondary production | G | +/- | +/- | (14, 19) |
|  | Horizontal  Biodiversity | Decomposition rate | H | +/- | +/- | (13, 14, 20) |
|  | Horizontal  Biodiversity | N loss | I | +/- | +/- | (18, 20–22) |
| endogenous | Fungal secondary production | N loss | J | 0 | - | (23–25) |
|  | Fungal secondary production | Decomposition rate | K | + | + | (23, 24) |
|  | N loss | Decomposition rate | L | + | + | (24, 26) |

**Table S2**: Site characteristic from measurements at the start and end of field experiment averaged across the 16 sites in Switzerland and Brazil. The numbers represent the mean ± standard deviation. b.d. indicates “below detection limit”, which was 0.07 µg L^-1^ for PO_4_^3-^ and 2.00 mgL^-1^ for DOC.

| Variable | Switzerland | | Brazil | |
| --- | --- | --- | --- | --- |
|  | forested | non-forested | forested | non-forested |
| Temperature [°C] | 4.21±1.15 | 4.41±1.40 | 20.86±1.75 | 21.15±1.84 |
| Velocity [ms^-1^] | 0.19±0.05 | 0.24±0.11 | 0.19±0.06 | 0.19±0.06 |
| pH | 7.58±0.51 | 7.56±0.46 | 6.76±0.11 | 6.68±0.32 |
| Conductivity [µScm^-1^] | 190.00±143.51 | 196.75±143.54 | 49.00±10.46 | 51.06±10.12 |
| NO_3_^-^ [mgL^-1^**]** | 2.44±2.19 | 2.22±2.20 | 0.76±0.29 | 0.79±0.32 |
| PO_4_^3-^ [µg/L | 7.87±1.95 | 3.82±8.62 | b.d. | b.d. |
| DOC [mgL^-1^] | 2.65±2.07 | 2.74±2.15 | b.d. | b.d. |
| O_2_ [%] | 98.94±1.41 | 99.29±2.54 | 106.93±11.61 | 102.41±8.87 |
| Alkalinity | 1.80±1.44 | 1.93±1.43 | 11.13±3.00 | 10.82±2.95 |
| Hardness | 1.16±0.83 | 1.22±0.82 | 24.38±10.84 | 26.38±8.12 |

**Table S3**: Initial leaf characteristics. Mean ± standard deviation.

| Taxon | %N | %C | %P | % fibre | % lignin |
| --- | --- | --- | --- | --- | --- |
| *Alnus* | 2.15±0.13 | 48.58±0.78 | 0.07±0.007 | 39.73±3.37 | 25.83±4.19 |
| *Fraxinus* | 1.52±0.23 | 42.71±4.64 | 0.09±0.007 | 36.59±3.52 | 19.57±5.04 |
| *Inga* | 0.90±0.28 | 33.32±8.38 | 0.02±0.001 | 76.31±0.91 | 49.39±0.75 |
| *Miconia* | 1.24±0.21 | 37.45±5.53 | 0.01±0.001 | 40.87±0.35 | 24.78±0.47 |

**Table S4**: Detailed model outputs for treatment effects on each function separately and as multifunctionality.

| Intercept_Slope | Estimate | EstError | l-95 | u-95 | Rhat | Bulk_ESS | Tail_ESS | Taxon | Model |
| --- | --- | --- | --- | --- | --- | --- | --- | --- | --- |
| Intercept | 0.46 | 0.28 | -0.23 | 1.05 | 1.01 | 1193 | 982 | *Alnus* | meanFunction ~ Mesh + Vegetation + Mix |
| MeshCoarse | 0.10 | 0.01 | 0.08 | 0.12 | 1.00 | 3926 | 2505 | *Alnus* | meanFunction ~ Mesh + Vegetation + Mix |
| Vegetationforested | 0.03 | 0.01 | 0.01 | 0.04 | 1.00 | 3379 | 2559 | *Alnus* | meanFunction ~ Mesh + Vegetation + Mix |
| MonoMixMix | -0.02 | 0.01 | -0.04 | 0 | 1.00 | 3744 | 2326 | *Alnus* | meanFunction ~ Mesh + Vegetation + Mix |
| Intercept | 0.80 | 0.27 | 0.19 | 1.39 | 1.02 | 189 | 134 | *Alnus* | biomass ~ Mesh + Vegetation + Mix |
| MeshCoarse | 0.06 | 0.01 | 0.05 | 0.07 | 1.00 | 1103 | 2554 | *Alnus* | biomass ~ Mesh + Vegetation + Mix |
| Vegetationforested | -0.01 | 0.01 | -0.02 | 0 | 1.00 | 2503 | 2471 | *Alnus* | biomass ~ Mesh + Vegetation + Mix |
| MixMix | 0.00 | 0.01 | -0.02 | 0.01 | 1.00 | 2142 | 2102 | *Alnus* | biomass ~ Mesh + Vegetation + Mix |
| Intercept | 0.23 | 0.18 | -0.15 | 0.68 | 1.00 | 1339 | 1284 | *Alnus* | N loss ~ Mesh + Vegetation + Mix |
| MeshCoarse | 0.11 | 0.02 | 0.08 | 0.14 | 1.00 | 3275 | 2382 | *Alnus* | N loss ~ Mesh + Vegetation + Mix |
| Vegetationforested | 0.04 | 0.02 | 0.01 | 0.07 | 1.00 | 3067 | 2618 | *Alnus* | N loss ~ Mesh + Vegetation + Mix |
| MonoMixMix | -0.01 | 0.01 | -0.04 | 0.02 | 1.00 | 3542 | 2515 | *Alnus* | N loss ~ Mesh + Vegetation + Mix |
| Intercept | 0.36 | 0.23 | -0.16 | 0.89 | 1.01 | 1310 | 1094 | *Alnus* | k ~ Mesh + Vegetation + Mix |
| MeshCoarse | 0.14 | 0.01 | 0.12 | 0.17 | 1.00 | 3951 | 2088 | *Alnus* | k ~ Mesh + Vegetation + Mix |
| Vegetationforested | 0.05 | 0.01 | 0.02 | 0.07 | 1.00 | 3523 | 2517 | *Alnus* | k ~ Mesh + Vegetation + Mix |
| MixMix | -0.05 | 0.01 | -0.07 | -0.02 | 1.00 | 3703 | 2558 | *Alnus* | k ~ Mesh + Vegetation + Mix |
| Intercept | 0.41 | 0.26 | -0.10 | 0.99 | 1.01 | 880 | 537 | *Fraxinus* | meanFunction ~ Mesh + Vegetation + Mix |
| MeshCoarse | 0.12 | 0.01 | 0.10 | 0.14 | 1.00 | 3684 | 2575 | *Fraxinus* | meanFunction ~ Mesh + Vegetation + Mix |
| Vegetationforested | 0.03 | 0.01 | 0.01 | 0.05 | 1.00 | 3307 | 2372 | *Fraxinus* | meanFunction ~ Mesh + Vegetation + Mix |
| MonoMixMix | 0.00 | 0.01 | -0.02 | 0.02 | 1.00 | 3104 | 2763 | *Fraxinus* | meanFunction ~ Mesh + Vegetation + Mix |
| Intercept | 0.83 | 0.27 | 0.18 | 1.39 | 1.01 | 418 | 475 | *Fraxinus* | biomass ~ Mesh + Vegetation + Mix |
| MeshCoarse | 0.04 | 0.01 | 0.03 | 0.05 | 1.00 | 2123 | 2612 | *Fraxinus* | biomass ~ Mesh + Vegetation + Mix |
| Vegetationforested | -0.01 | 0.01 | -0.02 | 0 | 1.00 | 2575 | 2443 | *Fraxinus* | biomass ~ Mesh + Vegetation + Mix |
| MixMix | 0.00 | 0.01 | -0.01 | 0.01 | 1.00 | 2932 | 2496 | *Fraxinus* | biomass ~ Mesh + Vegetation + Mix |
| Intercept | 0.56 | 0.62 | -0.16 | 1.6 | 1.65 | 6 | 19 | *Fraxinus* | N loss ~ Mesh + Vegetation + Mix |
| MeshCoarse | 0.16 | 0.01 | 0.14 | 0.19 | 1.09 | 44 | 314 | *Fraxinus* | N loss ~ Mesh + Vegetation + Mix |
| Vegetationforested | 0.05 | 0.01 | 0.02 | 0.08 | 1.38 | 41 | 672 | *Fraxinus* | N loss ~ Mesh + Vegetation + Mix |
| MonoMixMix | 0.01 | 0.01 | -0.02 | 0.04 | 1.41 | 76 | 826 | *Fraxinus* | N loss ~ Mesh + Vegetation + Mix |
| Intercept | 0.29 | 0.35 | -0.47 | 0.98 | 1.01 | 817 | 602 | *Fraxinus* | k ~ Mesh + Vegetation + Mix |
| MeshCoarse | 0.20 | 0.02 | 0.16 | 0.23 | 1.00 | 3603 | 2473 | *Fraxinus* | k ~ Mesh + Vegetation + Mix |
| Vegetationforested | 0.04 | 0.02 | 0.01 | 0.08 | 1.00 | 4147 | 2617 | *Fraxinus* | k ~ Mesh + Vegetation + Mix |
| MixMix | -0.02 | 0.02 | -0.05 | 0.02 | 1.00 | 4146 | 2925 | *Fraxinus* | k ~ Mesh + Vegetation + Mix |
| Intercept | 0.46 | 0.19 | 0.01 | 0.9 | 1.01 | 1035 | 813 | *Inga* | meanFunction ~ Mesh + Vegetation + Mix |
| MeshCoarse | 0.00 | 0.01 | -0.02 | 0.01 | 1.00 | 3178 | 2060 | *Inga* | meanFunction ~ Mesh + Vegetation + Mix |
| Vegetationforested | 0.02 | 0.01 | 0.00 | 0.03 | 1.00 | 3504 | 2263 | *Inga* | meanFunction ~ Mesh + Vegetation + Mix |
| MonoMixMix | 0.01 | 0.01 | -0.01 | 0.02 | 1.00 | 2923 | 2412 | *Inga* | meanFunction ~ Mesh + Vegetation + Mix |
| Intercept | 0.80 | 0.32 | -0.03 | 1.49 | 1.01 | 666 | 819 | *Inga* | biomass ~ Mesh + Vegetation + Mix |
| MeshCoarse | 0.00 | 0.01 | -0.02 | 0.02 | 1.00 | 1596 | 2172 | *Inga* | biomass ~ Mesh + Vegetation + Mix |
| Vegetationforested | 0.01 | 0.01 | -0.01 | 0.03 | 1.00 | 1560 | 2061 | *Inga* | biomass ~ Mesh + Vegetation + Mix |
| MixMix | -0.04 | 0.01 | -0.06 | -0.02 | 1.00 | 1504 | 2173 | *Inga* | biomass ~ Mesh + Vegetation + Mix |
| Intercept | 0.27 | 0.20 | -0.15 | 0.75 | 1.01 | 589 | 412 | *Inga* | N loss ~ Mesh + Vegetation + Mix |
| MeshCoarse | -0.01 | 0.01 | -0.04 | 0.02 | 1.00 | 3232 | 2574 | *Inga* | N loss ~ Mesh + Vegetation + Mix |
| Vegetationforested | 0.02 | 0.01 | 0.00 | 0.05 | 1.00 | 3300 | 2935 | *Inga* | N loss ~ Mesh + Vegetation + Mix |
| MonoMixMix | 0.05 | 0.01 | 0.02 | 0.08 | 1.00 | 2404 | 2403 | *Inga* | N loss ~ Mesh + Vegetation + Mix |
| Intercept | 0.42 | 0.52 | -0.44 | 2.02 | 1.05 | 86 | 45 | *Inga* | k ~ Mesh + Vegetation + Mix |
| MeshCoarse | 0.01 | 0.01 | -0.01 | 0.02 | 1.00 | 580 | 852 | *Inga* | k ~ Mesh + Vegetation + Mix |
| Vegetationforested | 0.02 | 0.01 | 0.00 | 0.04 | 1.01 | 579 | 749 | *Inga* | k ~ Mesh + Vegetation + Mix |
| MixMix | 0.01 | 0.01 | 0.00 | 0.03 | 1.01 | 604 | 680 | *Inga* | k ~ Mesh + Vegetation + Mix |
| Intercept | 0.46 | 0.24 | -0.16 | 1.03 | 1.02 | 1243 | 888 | *Miconia* | meanFunction ~ Mesh + Vegetation + Mix |
| MeshCoarse | 0.12 | 0.01 | 0.10 | 0.14 | 1.00 | 3858 | 2633 | *Miconia* | meanFunction ~ Mesh + Vegetation + Mix |
| Vegetationforested | 0.05 | 0.01 | 0.02 | 0.07 | 1.00 | 4075 | 2615 | *Miconia* | meanFunction ~ Mesh + Vegetation + Mix |
| MonoMixMix | 0.02 | 0.01 | 0.00 | 0.04 | 1.00 | 3861 | 2730 | *Miconia* | meanFunction ~ Mesh + Vegetation + Mix |
| Intercept | 0.68 | 0.27 | 0.04 | 1.26 | 1.01 | 1297 | 1127 | *Miconia* | biomass ~ Mesh + Vegetation + Mix |
| MeshCoarse | 0.02 | 0.01 | 0.00 | 0.04 | 1.00 | 4404 | 2588 | *Miconia* | biomass ~ Mesh + Vegetation + Mix |
| Vegetationforested | 0.02 | 0.01 | 0.00 | 0.04 | 1.00 | 3688 | 2694 | *Miconia* | biomass ~ Mesh + Vegetation + Mix |
| MixMix | 0.09 | 0.01 | 0.07 | 0.11 | 1.00 | 4100 | 2927 | *Miconia* | biomass ~ Mesh + Vegetation + Mix |
| Intercept | 0.32 | 0.27 | -0.26 | 0.89 | 1.01 | 1317 | 990 | *Miconia* | N loss ~ Mesh + Vegetation + Mix |
| MeshCoarse | 0.20 | 0.02 | 0.16 | 0.23 | 1.00 | 3271 | 2289 | *Miconia* | N loss ~ Mesh + Vegetation + Mix |
| Vegetationforested | 0.07 | 0.02 | 0.04 | 0.11 | 1.00 | 3478 | 2104 | *Miconia* | N loss ~ Mesh + Vegetation + Mix |
| MonoMixMix | -0.02 | 0.02 | -0.05 | 0.02 | 1.00 | 3278 | 2412 | *Miconia* | N loss ~ Mesh + Vegetation + Mix |
| Intercept | 0.36 | 0.29 | -0.32 | 1.04 | 1.01 | 1032 | 860 | *Miconia* | k ~ Mesh + Vegetation + Mix |
| MeshCoarse | 0.15 | 0.02 | 0.12 | 0.18 | 1.00 | 3507 | 2216 | *Miconia* | k ~ Mesh + Vegetation + Mix |
| Vegetationforested | 0.05 | 0.02 | 0.02 | 0.08 | 1.00 | 3428 | 2080 | *Miconia* | k ~ Mesh + Vegetation + Mix |
| MixMix | -0.02 | 0.01 | -0.04 | 0.01 | 1.00 | 4037 | 2164 | *Miconia* | k ~ Mesh + Vegetation + Mix |

**Table S5**: Detailed model outputs for each ecosystem function among themselves.

| Intercept_Slope | Estimate | EstError | l-95 | u-95 | Rhat | Bulk_ESS | Tail_ESS | Taxon | Model |
| --- | --- | --- | --- | --- | --- | --- | --- | --- | --- |
| Intercept | 0.79 | 0.27 | 0.08 | 1.37 | 1.01 | 619 | 368 | *Alnus* | Biomass ~ N loss |
| N_Loss.std | 0.06 | 0.03 | 0.00 | 0.12 | 1.00 | 2542 | 1207 | *Alnus* | Biomass ~ N loss |
| Intercept | 0.84 | 0.22 | 0.32 | 1.28 | 1.02 | 691 | 615 | *Fraxinus* | Biomass ~ N loss |
| N_Loss.std | 0.03 | 0.02 | -0.01 | 0.06 | 1.00 | 2894 | 2270 | *Fraxinus* | Biomass ~ N loss |
| Intercept | 0.82 | 0.37 | -0.08 | 1.66 | 1.01 | 614 | 627 | *Inga* | Biomass ~ N loss |
| N_Loss.std | -0.05 | 0.05 | -0.14 | 0.04 | 1.00 | 1496 | 2096 | *Inga* | Biomass ~ N loss |
| Intercept | 0.70 | 0.23 | 0.16 | 1.21 | 1.00 | 849 | 686 | *Miconia* | Biomass ~ N loss |
| N_Loss.std | 0.08 | 0.03 | 0.02 | 0.14 | 1.00 | 2464 | 1087 | *Miconia* | Biomass ~ N loss |
| Intercept | 0.75 | 0.28 | 0.20 | 1.42 | 1.04 | 155 | 72 | *Alnus* | Biomass ~ k |
| logk.std | 0.15 | 0.03 | 0.10 | 0.21 | 1.02 | 489 | 892 | *Alnus* | Biomass ~ k |
| Intercept | 0.85 | 0.25 | 0.39 | 1.27 | 1.01 | 574 | 354 | *Fraxinus* | Biomass ~ k |
| logk.std | 0.07 | 0.02 | 0.04 | 0.11 | 1.00 | 1436 | 1581 | *Fraxinus* | Biomass ~ k |
| Intercept | 0.70 | 0.44 | -0.47 | 1.70 | 1.03 | 132 | 92 | *Inga* | Biomass ~ k |
| logk.std | 0.20 | 0.07 | 0.06 | 0.33 | 1.00 | 569 | 1135 | *Inga* | Biomass ~ k |
| Intercept | 0.67 | 0.27 | 0.04 | 1.29 | 1.01 | 1338 | 1040 | *Miconia* | Biomass ~ k |
| logk.std | 0.18 | 0.04 | 0.10 | 0.26 | 1.00 | 2250 | 2281 | *Miconia* | Biomass ~ k |
| Intercept | 0.52 | 0.61 | -0.41 | 1.47 | 1.51 | 7 | 27 | *Alnus* | k ~ N loss |
| N_Loss.std | 0.76 | 0.03 | 0.70 | 0.84 | 1.18 | 156 | 2105 | *Alnus* | k ~ N loss |
| Intercept | 0.11 | 0.51 | -0.90 | 1.48 | 1.13 | 22 | 44 | *Fraxinus* | k ~ N loss |
| N_Loss.std | 0.80 | 0.02 | 0.76 | 0.85 | 1.04 | 95 | 255 | *Fraxinus* | k ~ N loss |
| Intercept | 0.28 | 0.26 | -0.28 | 0.92 | 1.01 | 675 | 338 | *Inga* | k ~ N loss |
| N_Loss.std | 0.29 | 0.04 | 0.22 | 0.36 | 1.00 | 2134 | 1118 | *Inga* | k ~ N loss |
| Intercept | 0.16 | 0.22 | -0.29 | 0.68 | 1.02 | 792 | 835 | *Miconia* | k ~ N loss |
| N_Loss.std | 0.66 | 0.03 | 0.61 | 0.72 | 1.00 | 1904 | 2400 | *Miconia* | k ~ N loss |

**Table S6**: Detailed model outputs for SEMs.

| model | Estimate | Est.Error | l95 | u05 | Rhat | Bulk_ESS | Tail_ESS | Taxon | Letter |
| --- | --- | --- | --- | --- | --- | --- | --- | --- | --- |
| logk_Intercept | -0.13 | 0.55 | -1.28 | 1.02 | 1.00 | 11728 | 11295 | *Alnus* | Intercept |
| NLoss_Intercept | -0.43 | 0.51 | -1.52 | 0.68 | 1.00 | 12304 | 11648 | *Alnus* | Intercept |
| logFungalBiomass_Intercept | -0.28 | 0.58 | -1.46 | 0.92 | 1.00 | 13036 | 13025 | *Alnus* | Intercept |
| logk_Coarse | 0.34 | 0.24 | -0.13 | 0.82 | 1.00 | 8364 | 10368 | *Alnus* | E |
| logk_Mix | -0.23 | 0.07 | -0.37 | -0.10 | 1.00 | 16222 | 13189 | *Alnus* | H |
| logk_Forested | 0.16 | 0.12 | -0.07 | 0.41 | 1.00 | 6318 | 8846 | *Alnus* | B |
| logk_ logFungalBiomass | 0.25 | 0.24 | -0.22 | 0.73 | 1.00 | 6286 | 9017 | *Alnus* | K |
| logk_NLoss | 0.50 | 0.27 | -0.05 | 1.03 | 1.00 | 5600 | 7756 | *Alnus* | L |
| NLoss_MeshCoarse | 0.57 | 0.15 | 0.27 | 0.86 | 1.00 | 13499 | 13752 | *Alnus* | F |
| NLoss_Mix | -0.07 | 0.11 | -0.29 | 0.16 | 1.00 | 23490 | 14256 | *Alnus* | I |
| NLoss_Forested | 0.36 | 0.12 | 0.13 | 0.59 | 1.00 | 24614 | 14992 | *Alnus* | C |
| NLoss_ logFungalBiomass | 0.26 | 0.14 | -0.03 | 0.53 | 1.00 | 10706 | 10279 | *Alnus* | J |
| logFungalBiomass _Coarse | 0.73 | 0.08 | 0.59 | 0.88 | 1.00 | 27887 | 15759 | *Alnus* | D |
| logFungalBiomass _Mix | -0.06 | 0.08 | -0.20 | 0.09 | 1.00 | 23189 | 15281 | *Alnus* | G |
| logFungalBiomass _Forested | -0.14 | 0.07 | -0.29 | 0.00 | 1.00 | 26258 | 15529 | *Alnus* | A |
| logk_Intercept | -0.08 | 0.52 | -1.18 | 1.01 | 1.00 | 13213 | 12821 | *Fraxinus* | Intercept |
| NLoss_Intercept | -0.67 | 0.48 | -1.74 | 0.40 | 1.00 | 14286 | 11712 | *Fraxinus* | Intercept |
| logFungalBiomass_Intercept | -0.26 | 0.59 | -1.52 | 0.98 | 1.00 | 16003 | 13545 | *Fraxinus* | Intercept |
| logk_Coarse | 0.19 | 0.28 | -0.35 | 0.78 | 1.00 | 9189 | 11370 | *Fraxinus* | E |
| logk_Mix | -0.16 | 0.07 | -0.29 | -0.03 | 1.00 | 17803 | 13019 | *Fraxinus* | H |
| logk_Forested | 0.13 | 0.11 | -0.07 | 0.38 | 1.00 | 7610 | 10104 | *Fraxinus* | B |
| logk_ logFungalBiomass | 0.21 | 0.25 | -0.27 | 0.72 | 1.00 | 6834 | 10685 | *Fraxinus* | K |
| logk_NLoss | 0.61 | 0.26 | 0.06 | 1.08 | 1.00 | 7211 | 9576 | *Fraxinus* | L |
| NLoss_MeshCoarse | 0.96 | 0.16 | 0.62 | 1.26 | 1.00 | 11875 | 12414 | *Fraxinus* | F |
| NLoss_Mix | 0.06 | 0.11 | -0.15 | 0.27 | 1.00 | 23933 | 15307 | *Fraxinus* | I |
| NLoss_Forested | 0.33 | 0.11 | 0.11 | 0.56 | 1.00 | 22584 | 15511 | *Fraxinus* | C |
| NLoss_ logFungalBiomass | 0.12 | 0.21 | -0.27 | 0.56 | 1.00 | 9166 | 9776 | *Fraxinus* | J |
| logFungalBiomass _Coarse | 0.60 | 0.10 | 0.41 | 0.79 | 1.00 | 28958 | 15214 | *Fraxinus* | D |
| logFungalBiomass _Mix | 0.07 | 0.10 | -0.12 | 0.26 | 1.00 | 25437 | 15564 | *Fraxinus* | G |
| logFungalBiomass _Forested | -0.17 | 0.09 | -0.35 | 0.02 | 1.00 | 29177 | 15399 | *Fraxinus* | A |
| logk_Intercept | -0.22 | 0.53 | -1.32 | 0.91 | 1.00 | 11042 | 12024 | *Inga* | Intercept |
| NLoss_Intercept | -0.25 | 0.45 | -1.25 | 0.72 | 1.00 | 11841 | 11205 | *Inga* | Intercept |
| logFungalBiomass_Intercept | 0.18 | 0.53 | -0.95 | 1.28 | 1.00 | 11386 | 12178 | *Inga* | Intercept |
| logk_Coarse | 0.08 | 0.10 | -0.12 | 0.28 | 1.00 | 20795 | 14798 | *Inga* | E |
| logk_Mix | 0.20 | 0.20 | -0.20 | 0.60 | 1.00 | 6992 | 11405 | *Inga* | H |
| logk_Forested | 0.15 | 0.12 | -0.08 | 0.39 | 1.00 | 12007 | 12999 | *Inga* | B |
| logk_ logFungalBiomass | 0.33 | 0.32 | -0.32 | 0.95 | 1.00 | 6151 | 9769 | *Inga* | K |
| logk_NLoss | 0.23 | 0.33 | -0.46 | 0.89 | 1.00 | 6258 | 9339 | *Inga* | L |
| NLoss_MeshCoarse | -0.08 | 0.12 | -0.31 | 0.15 | 1.00 | 30820 | 16454 | *Inga* | F |
| NLoss_Mix | 0.38 | 0.15 | 0.09 | 0.68 | 1.00 | 14319 | 14014 | *Inga* | I |
| NLoss_Forested | 0.19 | 0.12 | -0.04 | 0.42 | 1.00 | 28382 | 15065 | *Inga* | C |
| NLoss_ logFungalBiomass | -0.03 | 0.23 | -0.50 | 0.41 | 1.00 | 8043 | 9482 | *Inga* | J |
| logFungalBiomass _Coarse | -0.01 | 0.11 | -0.22 | 0.20 | 1.00 | 30341 | 16554 | *Inga* | D |
| logFungalBiomass _Mix | -0.42 | 0.11 | -0.63 | -0.21 | 1.00 | 34384 | 15007 | *Inga* | G |
| logFungalBiomass _Forested | 0.06 | 0.11 | -0.15 | 0.27 | 1.00 | 30942 | 15076 | *Inga* | A |
| logk_Intercept | 0.12 | 0.50 | -0.97 | 1.19 | 1.00 | 11313 | 11347 | *Miconia* | Intercept |
| NLoss_Intercept | -0.44 | 0.51 | -1.52 | 0.66 | 1.00 | 12554 | 11039 | *Miconia* | Intercept |
| logFungalBiomass_Intercept | -0.57 | 0.53 | -1.68 | 0.54 | 1.00 | 13124 | 12419 | *Miconia* | Intercept |
| logk_Coarse | 0.15 | 0.22 | -0.27 | 0.59 | 1.00 | 7305 | 8825 | *Miconia* | E |
| logk_Mix | -0.36 | 0.19 | -0.73 | 0.03 | 1.00 | 6475 | 9890 | *Miconia* | H |
| logk_Forested | -0.04 | 0.10 | -0.24 | 0.17 | 1.00 | 9320 | 11466 | *Miconia* | B |
| logk_ logFungalBiomass | 0.40 | 0.23 | -0.06 | 0.83 | 1.00 | 5912 | 9280 | *Miconia* | K |
| logk_NLoss | 0.67 | 0.23 | 0.21 | 1.10 | 1.00 | 7082 | 8928 | *Miconia* | L |
| NLoss_MeshCoarse | 0.91 | 0.11 | 0.69 | 1.11 | 1.00 | 20482 | 13830 | *Miconia* | F |
| NLoss_Mix | -0.33 | 0.21 | -0.73 | 0.10 | 1.00 | 7632 | 10907 | *Miconia* | I |
| NLoss_Forested | 0.30 | 0.11 | 0.08 | 0.51 | 1.00 | 17570 | 14539 | *Miconia* | C |
| NLoss_ logFungalBiomass | 0.31 | 0.24 | -0.20 | 0.76 | 1.00 | 6687 | 9566 | *Miconia* | J |
| logFungalBiomass _Coarse | 0.15 | 0.09 | -0.03 | 0.34 | 1.00 | 30745 | 14508 | *Miconia* | D |
| logFungalBiomass _Mix | 0.80 | 0.09 | 0.62 | 0.99 | 1.00 | 30503 | 14538 | *Miconia* | G |
| logFungalBiomass _Forested | 0.19 | 0.09 | 0.01 | 0.37 | 1.00 | 29440 | 15710 | *Miconia* | A |

**Table S7**: Untransformed mean ± SD for each ecosystem function, treatment and all four leaf species.

| Function | Treatment | *Alnus* | *Fraxinus* | *Inga* | *Miconia* |
| --- | --- | --- | --- | --- | --- |
| Decomposition rates  [dd-1] | mono  non-forested  micro | 0.00386±  0.00079 | 0.00649±  0.00116 | 0.00019±  0.00004 | 0.00047±  0.00013 |
|  | mix  non-forested  micro | 0.00343±  0.00079 | 0.00631±  0.00148 | 0.00019±  0.00004 | 0.00052±  0.00018 |
|  | mono  non-forested  micro+macro | 0.00446±  0.00078 | 0.00762±  0.00127 | 0.00019±  0.00004 | 0.00086±  0.00063 |
|  | mix  non-forested  micro+macro | 0.00415±  0.00073 | 0.00767±  0.00152 | 0.00019±  0.00004 | 0.00072±  0.00054 |
|  | mono  forested  micro | 0.00364±  0.00061 | 0.00637±  0.00083 | 0.00019±  0.00003 | 0.00051±  0.00015 |
|  | mix  forested  micro | 0.00348±  0.00059 | 0.00624±  0.00099 | 0.00021±  0.00006 | 0.00048±  0.00016 |
|  | mono  forested  micro+macro | 0.00534±  0.00159 | 0.00956±  0.00350 | 0.00021±  0.00009 | 0.00116±  0.00079 |
|  | mix  forested  micro+macro | 0.00513±  0.00205 | 0.00916±  0.00381 | 0.00023±  0.00017 | 0.00101±  0.00063 |
| Fungal secondary production  [mg/g] | mono  non-forested  micro | 60.88±  25.20 | 80.11±  22.88 | 22.74±  6.86 | 18.13±  8.68 |
|  | mix  non-forested  micro | 62.27±  26.22 | 83.81±  19.12 | 18.31±  5.90 | 29.50±  8.73 |
|  | mono  non-forested  micro+macro | 92.60±  36.40 | 106.87±  28.38 | 22.59±  7.16 | 21.96±  10.96 |
|  | mix  non-forested  micro+macro | 86.29±  22.22 | 101.56±  20.75 | 19.12±  6.32 | 30.40±  11.40 |
|  | mono  forested  micro | 63.01±  24.10 | 80.64±  20.80 | 22.02±  6.83 | 22.36±  8.71 |
|  | mix  forested  micro | 60.73±  19.22 | 85.85±  23.81 | 21.15±  8.92 | 28.63±  8.23 |
|  | mono  forested  micro+macro | 76.40±  21.97 | 94.02±  25.71 | 22.45±  6.56 | 23.48±  9.52 |
|  | mix  forested  micro+macro | 75.77±  27.52 | 94.22±  28.70 | 19.04±  5.85 | 30.30±  9.97 |
| N loss [%] | mono  non-forested  micro | 10.94±  5.01 | 16.08±  8.52 | -21.06±  10.02 | -2.04±  20.79 |
|  | mix  non-forested  micro | 10.32±  7.12 | 14.87±  10.63 | -17.86±  11.12 | 0.75±  22.18 |
|  | mono  non-forested  micro+macro | 14.72±  5.44 | 23.67±  8.58 | -23.98±  7.90 | 25.62±  31.77 |
|  | mix  non-forested  micro+macro | 14.49±  6.04 | 27.73±  12.71 | -16.46±  7.05 | 21.05±  25.13 |
|  | mono  forested  micro | 10.62±  3.92 | 13.25±  6.27 | -19.35±  11.32 | 7.47±  13.67 |
|  | mix  forested  micro | 8.96±  5.55 | 15.56±  6.02 | -14.10±  12.59 | 4.49±  16.41 |
|  | mono  forested  micro+macro | 22.87±  14.34 | 36.85±  20.37 | -20.25±  15.11 | 40.53±  28.11 |
|  | mix  forested  micro+macro | 21.60±  16.51 | 36.23±  18.20 | -15.72±  16.58 | 35.33±  28.64 |

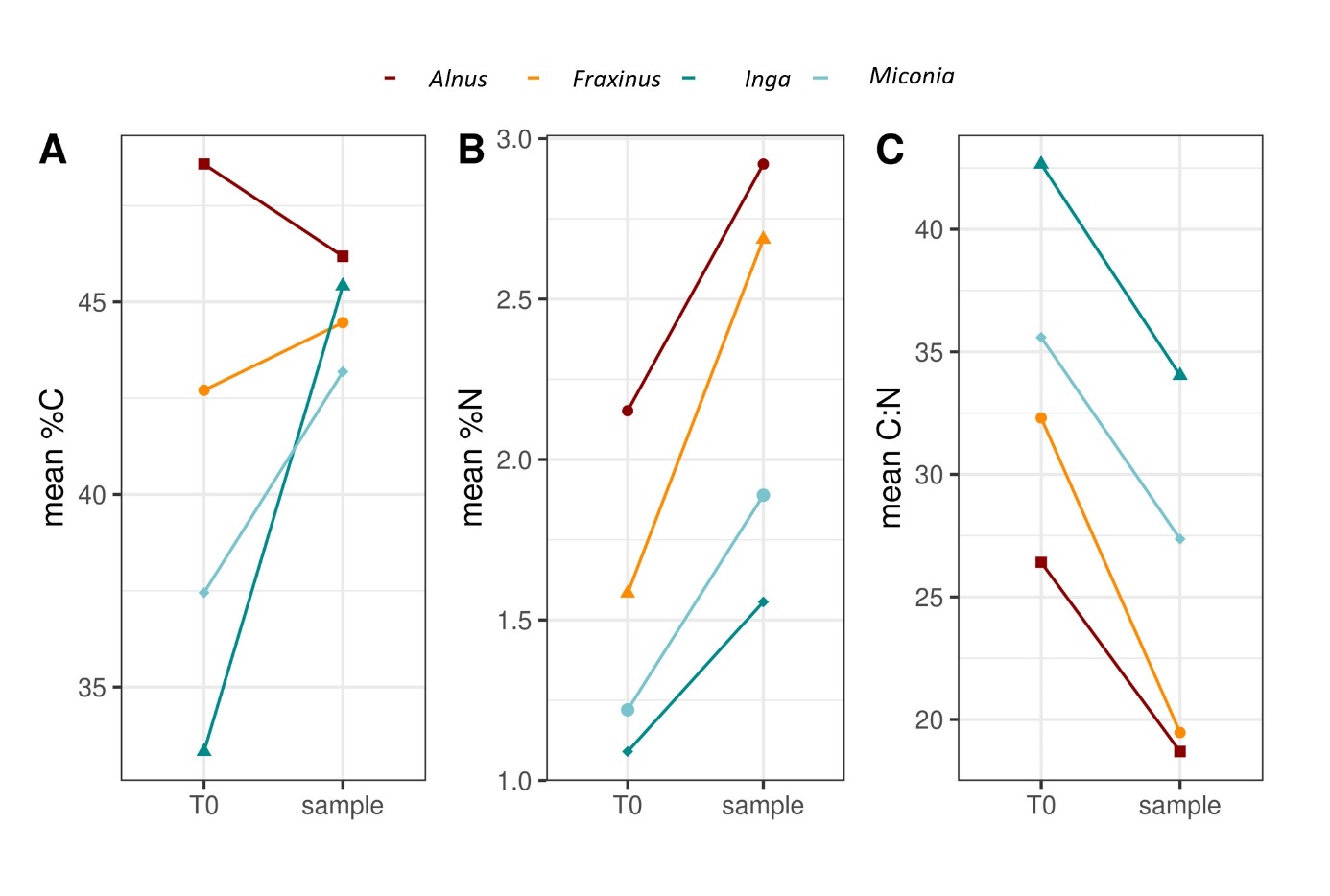

**Figure S1**: mean initial and experimental %C, %N and molar C:N ratios of leaf litter for the four leaf species *Alnus*, *Fraxinus*, *Inga* and *Miconia*.

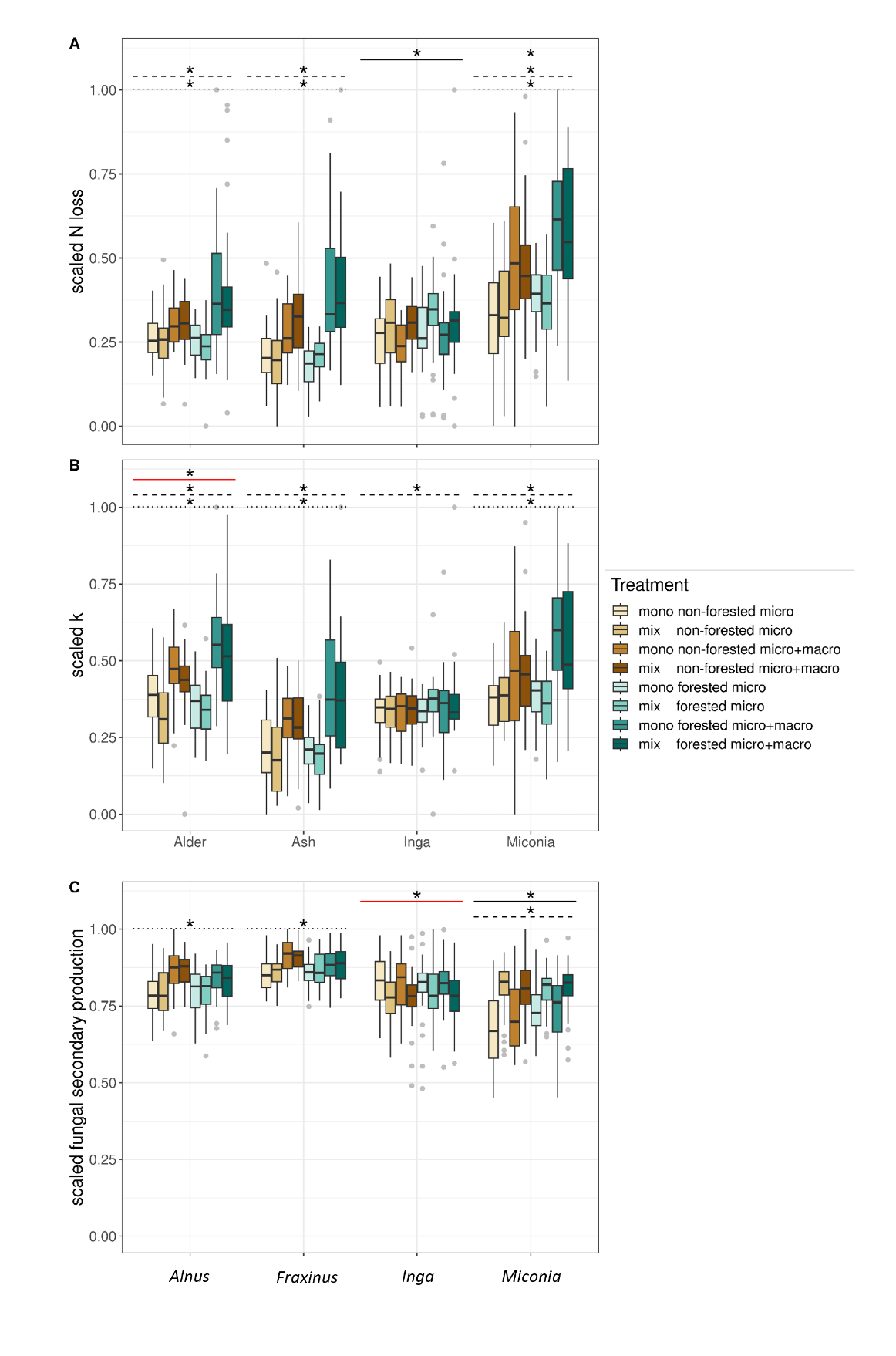

**Figure S2**: Boxplots of scaled mean multifunctionality depending on leaf species and experimental treatment. Colours differentiate the treatment combination of riparian vegetation type and horizontal and vertical biodiversity, with mixed (mix) or single (mono) species leaf litter bags, placement in a forested or non-forested stream section and inclusion (micro+macro) or exclusion (micro) of higher trophic level consumers. The solid lines represent statistically significant effects (0 not included in CIs) of horizontal biodiversity, dashed lines indicate effects of riparian vegetation type and dotted lines show effects of vertical biodiversity for each leaf species. Red coloured lines indicate that the effect was statistically significant in a negative direction.

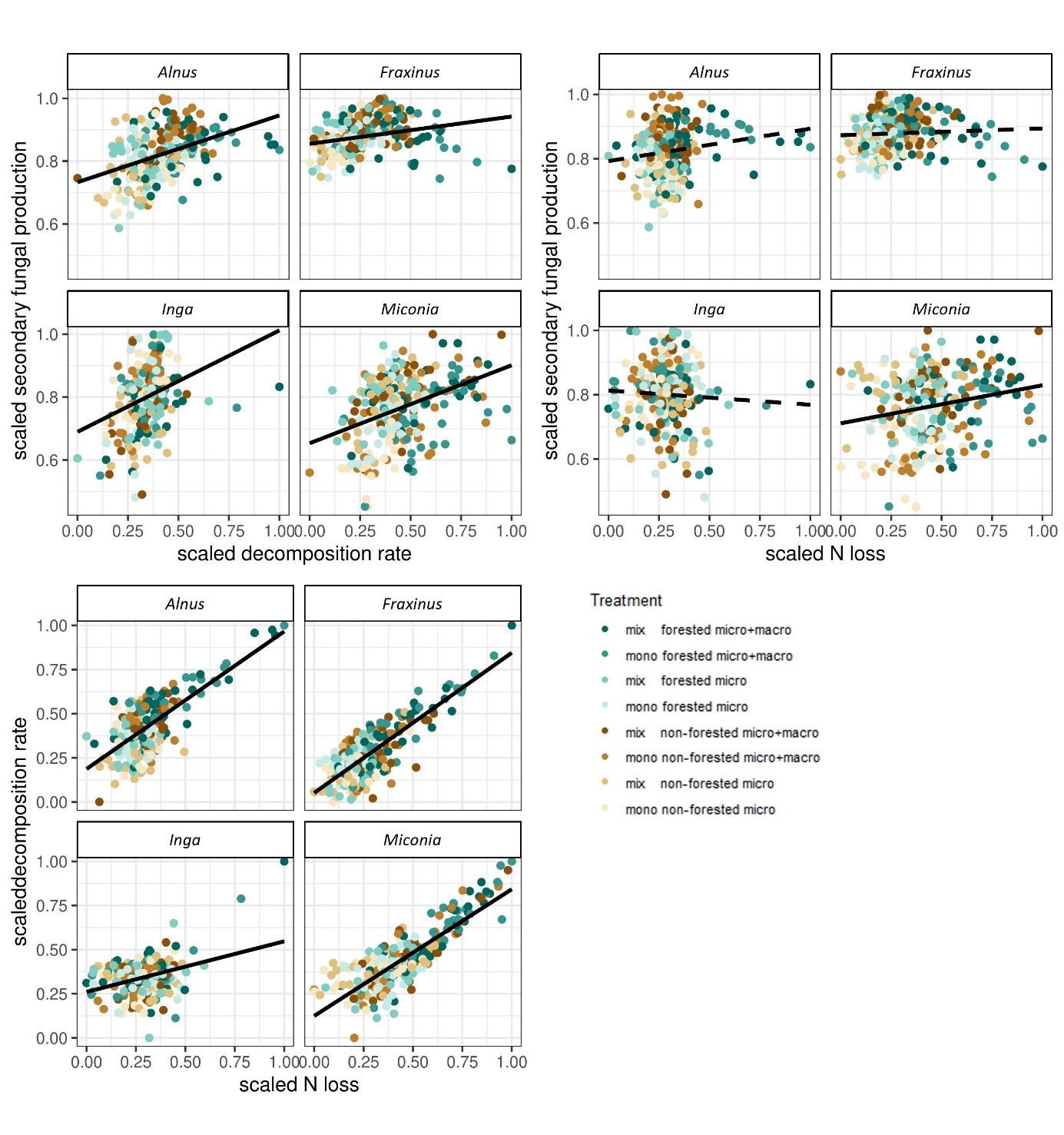

**Figure S3**: Relationships between ecosystem functions for the four leaf species. Solid lines indicate statistically significant slopes and the colours of the points show the different treatment combinations.
